# Myomerger induces fusion of non-fusogenic cells and is required for myoblast fusion

**DOI:** 10.1101/123158

**Authors:** Malgorzata E. Quinn, Qingnian Goh, Mitsutoshi Kurosaka, Dilani G. Gamage, Michael J. Petrany, Vikram Prasad, Douglas P. Millay

## Abstract

Despite the importance of cell fusion for mammalian development and physiology, the factors critical for this process remain to be fully defined^1^. This lack of knowledge has severely limited our ability to reconstitute cell fusion, which is necessary to decipher the biochemical mechanisms driving plasma membrane merger. Myomaker (*Tmem8c*) is a muscle-specific protein required for myoblast fusion^2,3^. Expression of myomaker in fibroblasts drives their fusion with myoblasts, but not with other myomaker-fibroblasts, highlighting the requirement of additional myoblast-derived factors for fusion. Here, we demonstrate that *Gm7325*, named myomerger, induces the fusion of myomaker-expressing fibroblasts. Cell mixing experiments reveal that while myomaker renders cells fusion-competent, myomerger induces fusogenicity. Thus, myomaker and myomerger confer fusogenic activity to normally non-fusogenic cells. Myomerger is skeletal muscle-specific and only expressed during developmental and regenerative myogenesis. Disruption of myomerger in myoblast cell lines through Cas9-mutagenesis generated non-fusogenic myocytes. Genetic deletion of myomerger in mice results in a paucity of muscle fibers demonstrating a requirement for myomerger in normal muscle formation. Myomerger deficient myocytes exhibit an ability to differentiate and harbor organized sarcomeres, however remain mono-nucleated. These data identify myomerger as a fundamental myoblast fusion protein and establishes a system that begins to reconstitute mammalian cell fusion.

---

The fusion of plasma membranes is necessary for numerous biological processes from conception to the development of skeletal muscle, osteoclasts, trophoblasts, and giant cells. The molecular regulation of fusion is poorly understood and the reconstitution of fusogenicity has not been achieved in mammalian cells. Specifically, the factors that directly participate in membrane coaelescence have not been identified. The development of systems that reconstitute fusion in the absence of other processes allow identification of nodal fusion machinery and associated molecular mechanisms. For example, the *C. elegans* fusogen epithelial fusion failure (Eff-1) is sufficient to fuse typically non-fusing cells^4,5^ and mechanisms of intracellular membrane fusion were partially revealed through reconstitution of SNAREs on synthetic membranes^6-8^. Thus, discoveries of specific fusion proteins and development of reconstitution systems have been historically critical to decipher multiple types of membrane fusion, however these systems are lacking for mammalian cellular fusion.

Myoblast fusion is a highly specific process essential for muscle formation, and while numerous proteins have been shown to regulate mammalian myoblast fusion^9-19^, myomaker is the only known muscle-specific protein absolutely required for this process^2^. Expression of myomaker in fibroblasts or mesenchymal stromal cells (MSCs) induces their fusion with muscle cells^20,21^. However, myomaker-fibroblasts do not fuse to each other indicating that additional muscle-derived factors are required for fusion. To uncover potential factors, we compared genes induced by expression of MyoD to their level of expression in 10T ½ fibroblasts. Of the top 100 MyoD-regulated genes not expressed in fibroblasts (Supplemental Information), we eliminated genes not likely to be directly involved in fusion (transcription factors, sarcomeric and metabolic genes) and focused on genes with transmembrane domains. This analysis yielded the following five genes: *Tmem182, Gm7325, Cdh15, Tspan33*, and *Tm6sf1*, however *Cdh15* was omitted from further analysis because it is not necessary for myoblast fusion or muscle formation^22^. We retrovirally expressed each gene in myomaker^+^ GFP^+^ fibroblasts and assayed for fusion. Appropriate expression in fibroblasts was verified through quantitative reverse transcription polymerase chain reaction (qRT-PCR) analysis (Extended Data Fig. 1a). We observed mainly mono-nucleated GFP^+^ cells in all cultures except when *Gm7325* was expressed where widespread multi-nucleated cells were present (Fig. 1a). Based on the ability of *Gm7325* to induce fusion of myomaker^+^ fibroblasts and the observations described below we named the protein myomerger.

**Figure 1.**
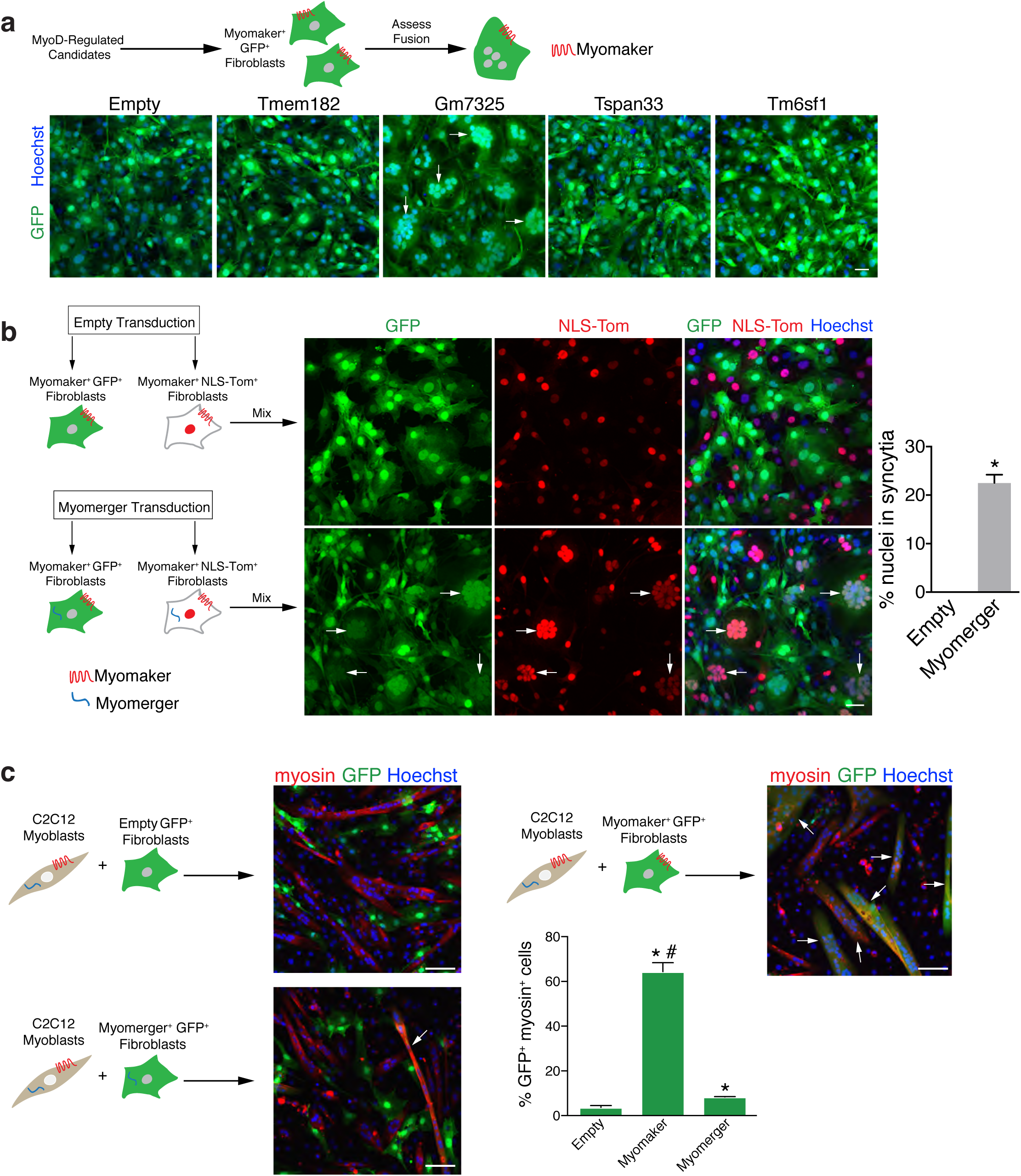
Induction of fibroblast fusion by myomerger. **a**, Schematic showing a screen for muscle genes that could activate fusion of GFP^+^ myomaker^+^ fibroblasts. Representative images of GFP^+^ cells and nuclei after expression of the indicated genes. Arrows depict cells with multiple nuclei. **b**, Illustration of cell mixing approach to show fusion between the populations of fibroblasts. Co-localization of GFP and NLS-TdTomato (NLS-Tom) in the nucleus represents fusion. Representative images demonstrate fusion of myomaker^+^ myomerger^+^ fibroblasts but not empty-infected myomaker^+^ fibroblasts. Arrows indicate fusion between GFP^+^ and NLS-Tom^+^ fibroblasts. The percentage of nuclei in syncytia after expression of empty or myomerger (*n*=3). **c**, Heterologous fusion experiment between C2C12 myoblasts and GFP^+^ fibroblasts infected with either empty, myomaker, or myomerger. Representative immunofluorescent images to visualize co-localization of myosin and GFP (arrows), indicating fusion. Quantification of the percentage of GFP^+^ myosin^+^ cells (*n*=3). Data are presented as mean ± SEM. \**P*<0.05 compared to empty. ^#^*P*<0.05 between myomaker and myomerger. Scale bars, 50 μm (**a, b**), 100 μm (**c**).

Multiple *Gm7325* transcripts are annotated in the University of California, Santa Cruz, mouse genome. The shorter transcript contains a single exon and yields a protein with 84 amino acids. In contrast, the longer transcript utilizes an upstream exon with an alternative start site and results in a protein of 108 amino acids (Extended Data Fig. 2a). The single coding exon of the short transcript is conserved in other mammalian genomes, including humans, while the upstream alternative exon leading to the longer transcript is not highly conserved (Extended Data Fig. 2b). For the initial screen we cloned the *Gm7325* locus into a retroviral vector, allowing normal splicing and expression of both short and long transcripts. Transduction of myomaker^+^ fibroblasts with either myomerger-short (S) or myomerger-long (L) induced formation of multi-nucleated cells, indicating both proteins are sufficient for fusion (Extended Data Fig. 2c). Additionally, myomerger and myomaker together induced fusion of 3T3 fibroblasts and MSCs (Extended Data Fig. 2d), suggesting these two genes could activate fusion in a multitude of cell types.

Given that multi-nucleated cells could arise from fusion or endoreplication, we designed a system to validate that multi-nucleated cells observed in fibroblasts expressing both myomerger and myomaker were generated through fusion. We engineered two fibroblast cell lines that both express myomaker, with one expressing GFP and the other expressing nuclear-localized TdTomato (NLS-Tom). Myomaker^+^ GFP^+^ and myomaker^+^ NLS-Tom^+^ fibroblasts were infected with a myomerger retrovirus or a control empty retrovirus, mixed, and fusion was assessed (Fig. 1b). We observed cells with multiple nuclei containing both GFP and NLS-Tom in fibroblasts expressing myomaker and myomerger indicating fusion (Fig. 1b). Quantification of fusion revealed approximately 20% of nuclei were contained in syncytia in cultures where fibroblasts were expressing both myomaker and myomerger (Fig. 1b). These results confirm that the observed syncytial cells are formed through fusion and that expression of myomaker and myomerger is sufficient to confer fusogenicity in non-fusogenic fibroblasts.

We also sought to determine the cell biology of fusion induced by myomaker and myomerger. We mixed myomaker^+^ myomerger^+^ GFP^+^ fibroblasts with NLS-Tom^+^ fibroblasts expressing myomaker or myomerger (Extended Data Fig. 3). Here we observed fusion of myomaker^+^ myomerger^+^ GFP^+^ fibroblasts with myomaker^+^ NLS-Tom^+^ but not myomerger^+^ NLS-Tom^+^ fibroblasts (Extended Data Fig. 3). This heterotypic nature of fibroblast fusion are consistent with our previously reported heterologous fusion system between myoblasts and fibroblasts^2^. In that system, myomaker^+^ fibroblasts that did not express myomerger fused with muscle cells, which express both myomaker and myomerger. We next utilized our heterologous fusion system where fibroblasts were infected with GFP and either empty, myomaker, or myomerger retrovirus, and then mixed with C2C12 myoblasts (Fig. 1c). In this assay, fusion is detected through co-localization of GFP (fibroblasts) with myosin^+^ myotubes. Compared to empty-infected GFP^+^ fibroblasts, we detected a significant increase in fusion between myosin^+^ cells with either myomaker^+^ GFP^+^ fibroblasts or myomerger^+^ GFP^+^ fibroblasts (Fig. 1c). However, quantification of myosin^+^ GFP^+^ cells revealed that myomerger did not drive the fusion of fibroblasts with muscle cells to the levels observed with myomaker (Fig. 1c). These data confirm myomaker is required in both fusing cells for *in vitro* fusion, while myomerger is essential in one fusing cell.

The only current information regarding *Gm7325* is its potential expression in embryonic stem (ES) cells^23^, therefore we interrogated its expression pattern more thoroughly. We performed qRT-PCR on multiple tissues from postnatal (P) day 5 mice with primers to distinguish the two potential mouse transcripts (Extended Data Fig. 4a). In neonatal tissues, we detected expression of both myomerger transcripts only in skeletal muscle (Fig. 2a). Despite the evidence of two myomerger transcripts, immunoblot analysis of skeletal muscle lysates from P5 mice using a commercially available antibody identified a single band at approximately 12 kDa. This band was absent in P28 lysates indicating that myomerger is downregulated after neonatal development (Fig. 2b). Skeletal muscle exhibits a robust ability to regenerate due to the presence of muscle stem cells, also known as satellite cells^24,25^. We analyzed expression of myomerger in *mdx*^4cv^ mice, which is a mouse model of muscular dystrophy that leads to chronic cycles of degeneration and regeneration^26^. Myomerger expression was detected in diaphragm lysates from *mdx*^4cv^ mice, but not control diaphragms (Fig. 2c). Finally, we analyzed expression of myomerger in a model of skeletal muscle overload-induced (MOV) hypertrophy and observed up-regulation (Fig. 2d). Collectively, these data demonstrate that myomerger is expressed only during development and is induced upon satellite cell activation in the adult.

**Figure 2.**
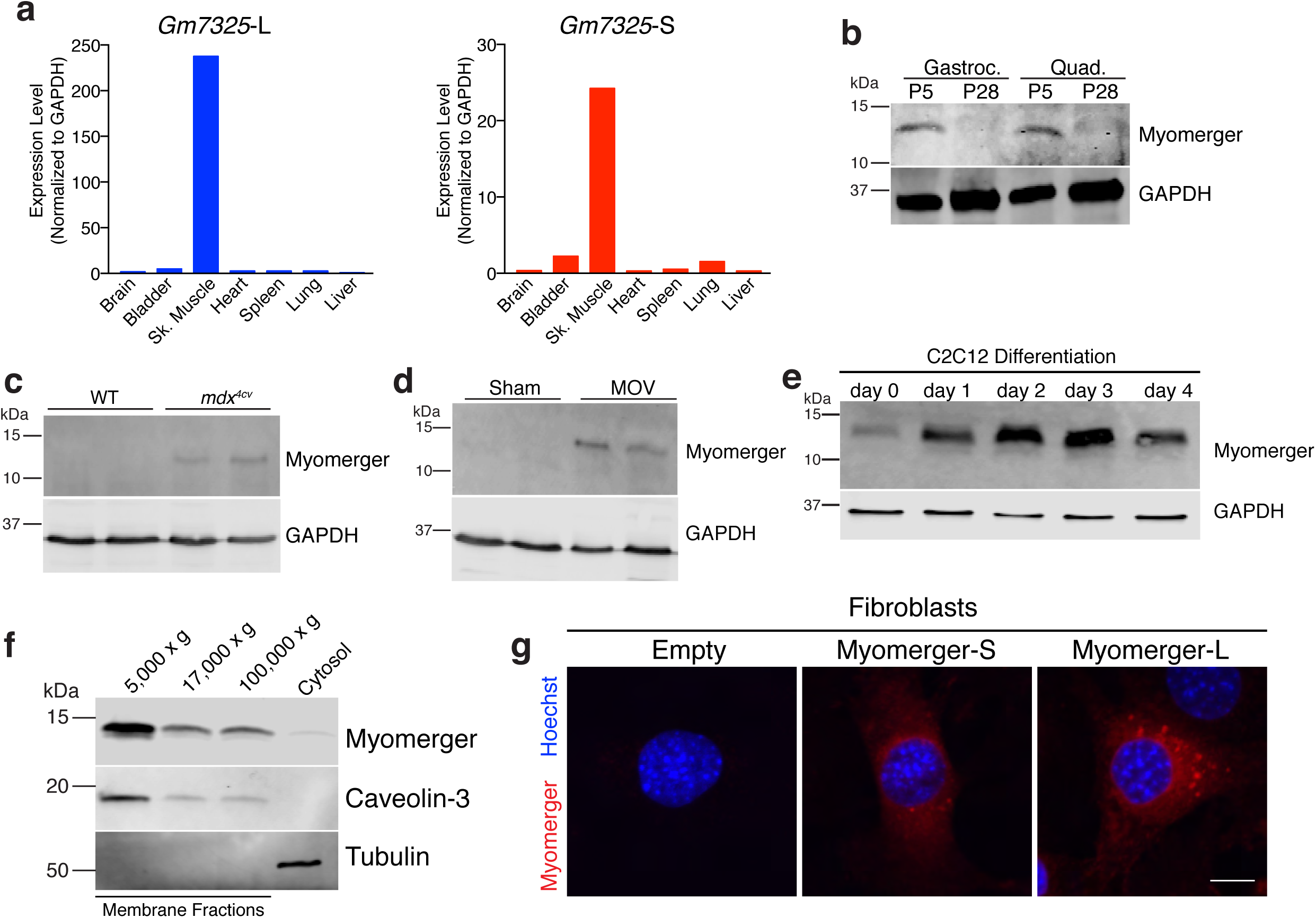
Muscle-specific expression and regulation of myomerger. **a**, qRT-PCR for both *Gm7325* long (L) and short (S) transcripts from various postnatal (P) day 5 tissues. **b-e**, Immunoblotting for myomerger comparing P5 muscle to P28 muscle (**b**), WT to *mdx^4cv^* diaphragms (**c**), sham plantaris to mechanically overloaded (MOV) plantaris (**d**), and during C2C12 myoblast differentiation (**e**). GAPDH was used as a loading control. **f**, Fractionation of C2C12 lysates on day 2 of differentiation followed by immunoblotting. **g**, Representative immunostaining of fibroblasts infected with either empty, myomerger-short (S), or myomerger-long (L). Scale bar, 10 μm.

We next sought to determine if myomerger is regulated as myoblasts differentiate. In C2C12 cells, both *Gm735* transcripts were significantly elevated during differentiation (Extended Data Fig. 4b). Similarly, myomerger protein levels were low in proliferating myoblasts (day 0), but increased upon differentiation with expression maintained during myoblast differentiation and fusion into myotubes (Fig. 2e). The short mouse myomerger protein, but not the long form, is highly conserved among vertebrate species (Extended Data Fig. 4c). After transduction of C2C12 cells with empty, myomerger-S, or myomerger-L we detected an increased upper band in cells expressing either myomerger-S or myomerger-L that co-migrated with the 12 kDa endogenous protein in the empty-infected C2C12 cells (Extended Data Fig. 4d). A lower band was identified exclusively in myomerger-S lysates suggesting that complex mRNA or post-translational processing results in the endogenous single 12 kDa band observed in WT C2C12 cells and muscle homogenates. Both myomerger proteins harbor a hydrophobic region close to the N-terminus, where computational analysis of this region indicates a signal peptide or transmembrane domain (Extended Data Fig. 4e). Given that both variants were found to confer fusogenicity, the significance of the predicted domains is presently unclear. To understand subcellular localization, we fractionated C2C12 cells on day 2 of differentiation and identified myomerger in membrane fractions containing caveolin-3, a protein known to associate with both heavy and light vesicles (Fig. 2f). Immunostaining of fibroblasts expressing myomerger-S or myomerger-L shows that both proteins exhibit similar diffuse and vesicular localization (Fig. 2g). Thus, myomerger associates with membrane compartments consistent with its ability to induce fusion.

The ability of myomerger to induce fusion of myomaker-fibroblasts, and its muscle-specific expression in the mouse, suggests that myomerger may play a critical role during myogenesis. To begin to decipher the role of myomerger in myogenesis, we evaluated its function during myoblast differentiation. We utilized CRISPR/Cas9 genome editing to disrupt myomerger in C2C12 myoblasts. Two guide RNAs (gRNA) were designed to target the largest exon of *Gm7325*, which resulted in a 166 base pair deletion thereby disrupting both mouse transcripts (Extended Data Fig. 5a). C2C12 cells were transfected with a plasmid containing Cas9 with an IRES-GFP and myomerger gRNAs, or transfected with only Cas9-IRES-GFP as a control. Flow cytometry of GFP^+^ cells followed by genotyping through PCR analysis revealed disruption of the myomerger locus (Extended Data Fig. 5b). Myomerger was not detectable in myomerger KO C2C12 cells confirming efficient disruption of the locus (Fig. 3a). Control and myomerger KO C2C12 cells were then analyzed for their ability to differentiate and form myotubes. WT myoblasts differentiated, as indicated by myosin^+^ cells, and fused to form multi-nucleated myotubes (Fig. 3b). In contrast, myomerger KO C2C12 cells exhibited the ability to differentiate but lacked fusogenic activity to form myotubes (Fig. 3b). Indeed, quantification of the differentiation index revealed no difference in the percentage of myosin^+^ cells between WT and myomerger KO cultures (Fig. 3c). Additionally, quantification of fusion demonstrated that myomerger KO myosin^+^ cells remain mono-nucleated while WT cells fuse (Fig. 3d). qRT-PCR analysis for the myogenic genes *Myogenin, Myh4, Ckm*, and *Tmem8c* (myomaker) further indicated that myomerger KO myoblasts activate the differentiation program (Fig. 3e). Interestingly, myogenic transcripts were elevated in myomerger KO cells potentially suggesting a feedback mechanism by which non-fusogenic cells attempt to further differentiate (Fig. 3e). Infection of myomerger KO C2C12 cells with either myomerger-S or myomerger-L rescued the fusion defect demonstrating that the phenotype in these cells is specifically due to the loss of myomerger (Extended Data Fig. 5c). Western blot analysis from these lysates shows re-expression of myomerger in KO cells (Extended Data Fig. 5d). As a potential mechanism for the lack of fusion in myomerger KO myocytes, we examined expression and localization of myomaker. On day 2 of differentiation, myomerger KO cells exhibited normal expression and localization of myomaker (Extended Data Fig. 6a). Moreover, we did not detect widespread co-localization between myomaker and myomerger suggesting that myomerger does not directly regulate myomaker distribution (Extended Data Fig. 6b). These data reveal that myomerger is necessary for myoblast fusion *in vitro*.

**Figure 3.**
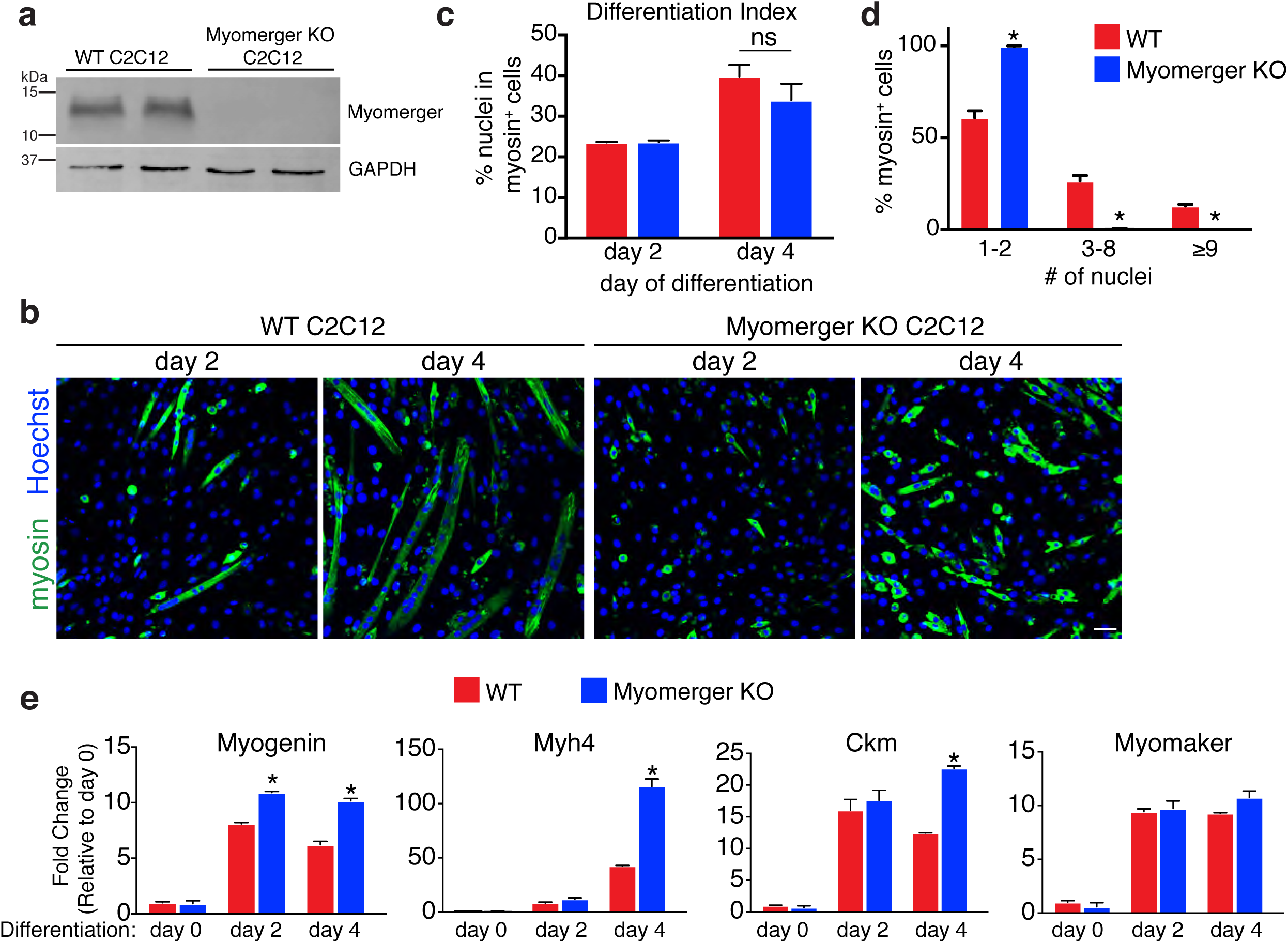
Requirement of myomerger for myoblast fusion *in vitro*. **a**, Immunoblotting for myomerger in WT and myomerger KO C2C12 cells on day 2 of differentiation. GAPDH was used as a loading control. **b**, Representative immunofluorescence images on day 2 and day 4 of differentiation for WT and myomerger KO C2C12 cells. Myomerger KO cells differentiate but fail to fuse. **c**, Quantification of the differentiation index, the percentage of nuclei in myosin^+^ cells (*n*=4). ns, not significant. **d**, The percentage of myosin^+^ cells that contain 1-2, 3-8, or ≥9 nuclei after 4 days of differentiation, as an indicator of fusogenicity (*n*=3). **e**, qRT-PCR for the indicated myogenic transcripts (*n*=4). Data are presented as mean ± SEM. \**P*<0.05 compared to WT. Scale bar, 50 μm.

To examine the function of myomerger *in vivo*, we disrupted exon 3 using the same CRISPR/Cas9 strategy described for C2C12 myoblasts. Injection of Cas9 and myomerger gRNAs into blastocysts resulted in lethality of 9 of the 10 F_0_ pups, suggesting that the high efficiency of Cas9 lead to homozygous deletion of myomerger. The one remaining pup was heterozygous for myomerger and was mated to generate *Gm7325^-/-^* mice (Extended Data Fig. 7a). Sequencing the mutant PCR product from the heterozygous founder revealed the presence of the same mutation as was achieved in C2C12 cells. We failed to observe any *Gm7325^-/-^* mice upon genotyping two litters at P7 suggesting that, in conjunction with the lethality detected through initial generation of F_0_ pups, myomerger is essential for life. Indeed, E17.5 *Gm7325^-/-^* embryos exhibited minimal skeletal muscle upon gross examination (Fig. 4a). Specifically, bones of the limbs and rib cage were noticeable due to a scarcity of surrounding muscle as observed in WT embryos. Myomerger KO mice also display a hunched appearance with elongated snouts, hallmark characteristics in embryos with improper muscle formation (Fig. 4a). Detection of myomerger by western blot of WT and *Gm7325^-/-^* tongues showed elimination of myomerger protein in KO samples (Extended Data Fig. 7b). E15.5 forelimb sections show that myomerger KO myoblasts express myogenin indicating that specification and differentiation are activated despite loss of myomerger (Fig. 4b). Moreover, histological analysis of multiple muscle groups at E15.5 revealed the presence of myosin^+^ muscle cells and sarcomeric structures in myomerger KO mice, (Fig. 4c and Extended Data Fig. 7c). While multi-nucleated myofibers were present in WT mice, these structures were not readily detected in myomerger KO mice indicating that genetic loss of myomerger renders myocytes non-fusogenic (Fig. 4c and Extended Data Fig. 7c). Analysis of forelimbs from E17.5 WT and myomerger KO embryos confirm that myomerger KO myoblasts are unable to properly fuse, although we did detect myocytes with two nuclei (Extended Data Fig. 7d-f). These results, together with our *in vitro* analysis, reveals that myomerger is required for muscle formation during mammalian development through regulation of myoblast fusion.

**Figure 4.**
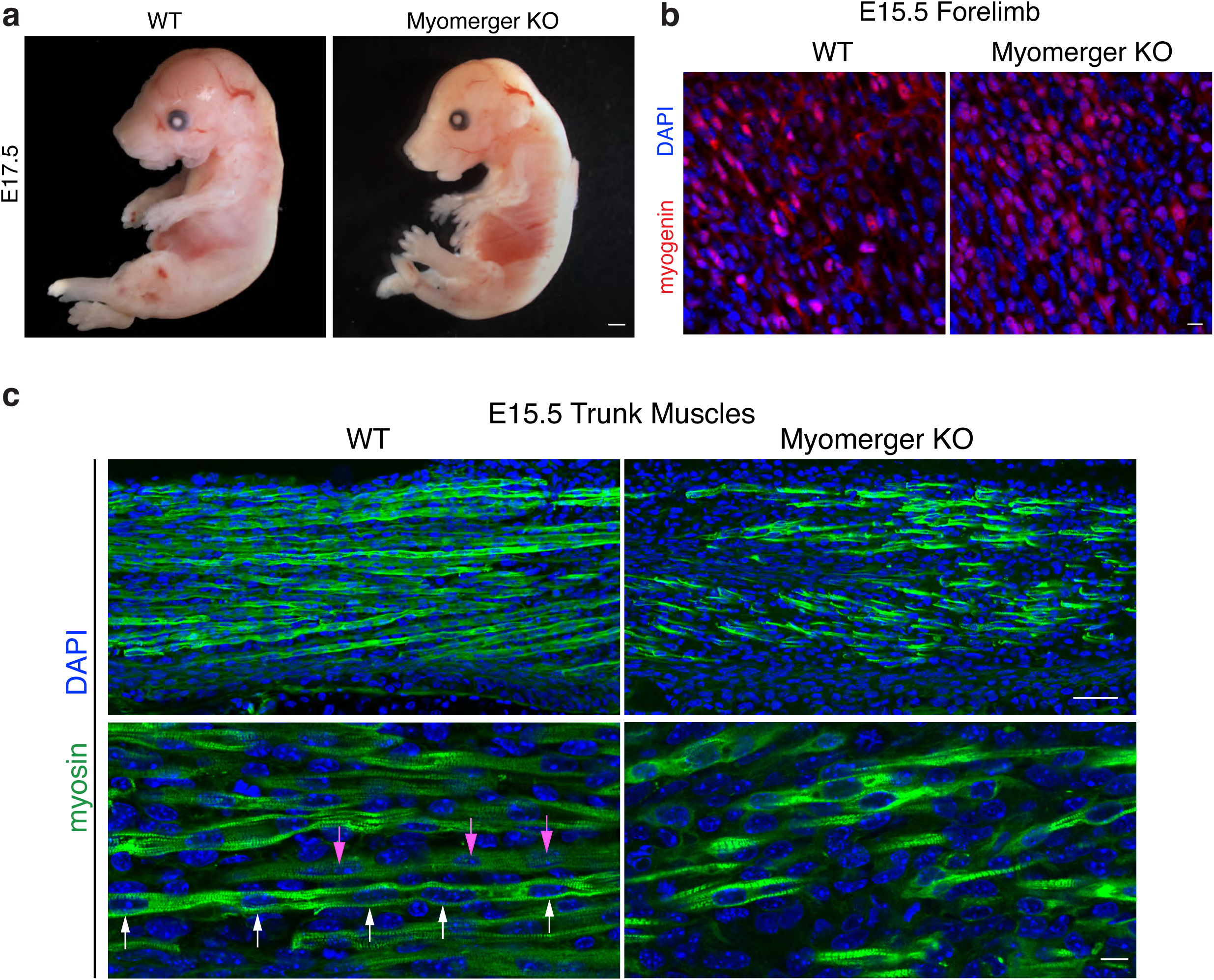
Myomerger is essential for myoblast fusion and muscle formation during embryonic development. **a**, Representative whole-mount images of WT and myomerger KO E17.5 embryos showing improper skeletal muscle formation in KO embryos (*n*=4). **b**, Immunofluorescence images for myogenin from WT and myomerger KO E15.5 forelimbs demonstrating that myomerger is not required for myogenic activation (*n*=3). **c**, Myosin immunofluorescence on the indicated E15.5 trunk muscles (*n*=3). Multi-nucleated myofibers (arrows of same color show nuclei within one myofiber) were observed in WT sections. Myomerger KO myocytes were myosin^+^ with sarcomeres but remained mono-nucleated. Scale bars, 1 mm (**a**), 50 μm (**c**, top panels), 10 μm (**b, c**, bottom panels).

In summary, we report the discovery of an additional muscle-specific factor required for myoblast fusion and developmental myogenesis. While myomaker and myomerger are both necessary for muscle formation, E17.5 myomerger KO embryos exhibit more myocytes compared to embryos lacking myomaker suggesting that these two key myoblast fusion proteins may have distinct functions. Given that myomaker and myomerger are both essential for myoblast fusion, an understanding of their precise function will aid in the delineation of mammalian plasma membrane fusion mechanisms. The fibroblast cell fusion system developed here, through expression of myomaker and myomerger, provides a unique platform to decipher fusion mechanisms. Moreover, the induction of cell fusion by two factors represents an avenue for gene delivery to potentially any tissue.

## Methods

### Cell Culture

C2C12 cells, 10T 1/2 fibroblasts, and NIH/3T3 fibroblasts were purchased from American Type Culture Collection and propagated in DMEM (Gibco) containing 10% heat-inactivated bovine growth serum (BGS) and supplemented with antibiotics. C2C12 cells were differentiated by switching to media containing 2% heat-inactivated horse serum (HS) and antibiotics. MSCs were a gift from Jose Cancelas and described previously^27^.

### Bioinformatic Analysis

Microarray data from the GEO DataSet GSE34907^28^ was interrogated using GEO2R analysis to identify 1826 genes displaying an increase greater than 1 log fold-change in MyoD-expressing fibroblasts. In parallel, a transcriptional profile of 10T 1/2 fibroblasts transduced with empty virus was generated using RNA-seq analysis (paired-end library layout using Illumina sequencing platform) and a list of all genes with RPKM values below 1.5 compiled using Strand NGS software (Ver. 2.6; Build: Mouse mm10 (UCSC) using Ensembl transcript annotations). These two gene lists were then compared to generate a final tally comprised of 531 genes that were both upregulated in MyoD-expressing fibroblasts and had low or no detectable expression in 10T 1/2 fibroblasts. Finally, the top 100 genes were interrogated for genes that contain transmembrane domains and not previously studied for their role during myoblast fusion.

### Animals

We used a dual sgRNA targeting strategy to create *Gm7325^-/-^* mice. We selected the sgRNAs according to the on- and off-target scores from the web tool CRISPOR^29^. The selected gRNAs were 5’-GCAGCGATCGAAGCACCATC-3’ and 5’-GAGGCCTCTCCAGAATCCGG-3’ that target exon 3 of *Gm7325*. The sgRNAs were *in vitro* synthesized using the MEGAshortscript T7 kit (ThermoFisher) and purified by the MEGAclear Kit (ThermoFisher), as previously described^30^. sgRNAs (50 ng/ul of each) were mixed with 100 ng/ul Cas9 protein (ThermoFisher) and incubated at 37°C for 15 min to form a ribonucleoprotein complex. We then injected the mix into the cytoplasm of one-cell-stage embryos of the C57BL/6 genetic background using a piezo-driven microinjection technique as described previously^30^. Injected embryos were immediately transferred into the oviducal ampulla of pseudopregnant CD-1 females. Live born pups were genotyped by PCR with the following primers: F: 5′- GAAGGGAGGACTCCACACCC-3’ and R: 5’-CGCCTGGACTAACCGGCTCC-3’. The edited allele was further confirmed by Sanger sequencing. One heterozygous founder was obtained and mated with WT C57Bl6 mice to eventually generate KO mice. *Mdx^4cv^* mice were purchased from Jackson Laboratory (#002378). Muscle overload of the plantaris muscle was achieved through bilateral synergistic ablation of soleus and gastrocnemius muscles as described by others^31^. Briefly, the soleus and gastrocnemius muscles were exposed by making an incision on the posterior-lateral aspect of the lower limb. The distal and proximal tendons of the soleus, lateral and medial gastrocnemius were subsequently cut and carefully excised. All animal procedures were approved by Cincinnati Children’s Hospital Medical Center’s Institutional Animal Care and Use Committee.

### CRISPR-Mediated Genome Editing in C2C12 Cells

Freshly plated low passage C2C12 cells were transfected with 4μg of a modified pX458 plasmid (Addgene #48138, gift from Yueh-Chiang Hu), which contained a high fidelity Cas9, an optimized sgRNA scaffold, and an IRES-GFP cassette. The same gRNAs used to generate KO animals were used for C2C12 cells. 16 μL of Lipofectamine 2000 was used for this transfection. 5 × 10^5^ C2C12 cells were transfected in a 60 mm culture dish. Forty-eight hours after transfection GFP^+^ cells were sorted into 96 well plates using FACS. These cells were maintained in DMEM containing 20% FBS with antibiotics at subconfluent densities. The cell lines were genotyped by amplifying a 420 bp region surrounding the site of Cas9 activity using the primers used to genotype *Gm7325^-/-^* animals.

### Cloning and viral infection

We initially cloned a region of the *Gm7325* locus, containing all genomic information for expression of myomerger-short and myomerger-long, from C57Bl6 mouse genomic DNA using the following primers: F: 5’-AGTGATGCTGAATCCACCGCA-3’ and R: 5’-CCAATAACAACACACTGTCCT-3’. We cloned myomerger-short and long coding sequences from cDNA of differentiating C2C12 cells using the following primers: myomerger-long F: 5’-ATGCCAGAAGAAAGCTGCACTG-3’, myomerger-short F: 5’- ATGCCCGTTCCATTGCTCCCGA-3’, and a common myomerger R: 5’-TCA CTT CTG GGG GCC CAA TCT C-3’. Myomerger cDNA and genomic DNA was cloned into the retroviral vector pBabe-X^2^ using EcoRI. Myomaker and GFP retroviral plasmids have been described previously^2^. NLS-TdTomato was subcloned from pQC-NLS-TdTomato (Addgene #37347) into the retroviral vector pMX (Cell Biolabs). Plasmids containing cDNA for Tmem182, Tspan33, and Tm6sf1 from the Mammalian Gene Collection were purchased from Open Biosystems and subcloned into pBabe-X. Ten micrograms of retroviral plasmid DNA were transfected with FuGENE 6 (Roche) into Platinum E cells (Cell Biolabs), which were plated 24 hours before transfection on a 10 cm culture dish at a density of 3-4×10^6^ cells per dish. Forty-eight hours after transfection, viral media were collected, filtered through a 0.45 μm cellulose syringe filter and mixed with polybrene (Sigma) at a final concentration of 6 μg/ml. Target cells were plated on 10 cm culture dishes at a density of 4×10^5^ cells per dish 16-18 hours before infection. Eighteen hours after infection, virus was removed, cells were washed with PBS and split into new 10 cm dishes.

### Cell fusion assays

Cells were split 18 hours after retroviral infection and split again 24-48 hours later. At the second split, cells were seeded for the fusion assay on 35-mm dishes (3-4×10^5^ cells per dish) or on 8- well Ibidi slides (2×10^4^ cells/well). Fusion was assessed 24-48 hours after seeding. For heterologous fusion, cultures of fibroblasts and myoblasts mixed at a ratio of 1:1 (1.5×10^5^ cells for each) were induced to differentiate 24 hours after seeding and fusion was assessed on day 4 of differentiation.

### RNA extraction and quantitative RT-PCR (qRT-PCR)

Total RNA was extracted from either mouse tissue or cultured cells with TRIZOL (Life Technologies) according to manufacturer’s protocol. cDNA was synthesized using High-Capacity cDNA Reverse Transcription Kit (Applied Biosystems) with random primers. Gene expression was assessed using standard quantitative PCR approach with Power Sybr^®^ Green PCR mastermix (Applied Biosystems). Analysis was performed on StepOnePlus Real-Time PCR system (Applied Biosystems) with the following primers: myomerger-short-F: 5′-CAGGAGGGCAAGAAGTTCAG-3′ and myomerger-R: 5′-ATGTCTTGGGAGCTCAGTCG-3′; myomerger-long-F: 5′-ACCAGCTTTCATGCCAGAAG-3′ and myomerger-R: 5′- ATGTCTTGGGAGCTCAGTCG-3′; myomaker-F: 5′-ATCGCTACCAAGAGGCGTT-3′ and myomaker-R: 5′-CACAGCACAGACAAACCAGG-3′; Tm6sf1 F: 5’-TTAGTGGTCCCTGGATGCTC-3” and Tm6sf1 R: 5’-GACGCACCAATGTGAGAAAA-3’; Tspan33-F: 5’-GGGGACGAGTTCTCCTTCG-3’ and Tspan33 R: 5’-TGCTTCTGCGTGCTTCATTAG-3’; Tmem182-F: 5’-GGCTCTCTTCGGAGCTTTGG-3’ and Tmem182-R: 5’-GGTGGCTGATTGGTGTACCAG-3’; myogenin-F: 5’-CTACAGGCCTTGCTCAGCTC-3’ and myogenin-R: 5’-GTGGGAGTTGCATTCACTGG-3’; Ckm-F: 5’-ACCTCCACAGCACAGACAGA-3’ and Ckm-R: 5’-CAGCTTGAACTTGTTGTGGG-3’; Myh4-F: 5’-GCAGGACTTGGTGGACAAAC-3’ and Myh4-R: 5’-ACTTGGCCAGGTTGACATTG-3’; GAPDH-F: 5’-TGCGACTTCAACAGCAACTC-3’ and GAPDH-R: 5’- GCCTCTCTTGCTCAGTGTCC-3’.

### Western blot analysis

Cultured cells were washed two times with ice cold PBS, scraped into a conical tube, pelleted, resuspended in lysis buffer (50 mM Tris-HCl, pH 6.8, 1mM EDTA, 2% SDS) and sonicated for a total of 15 seconds (three 5 second pulses). Skeletal muscle tissues from mice were homogenized with a bead homogenizer (TissueLyser II; Qiagen) in lysis buffer (10 mM Tris (pH 7.4), 1 mM EDTA, 1mM dithiothreitol, 0.5% Triton X-100, 2.1 mg/ml NaF) containing protease and phosphatase inhibitor cocktails (5 μl/ml; Sigma-Aldrich). Both cells and tissue lysates were centrifuged to pellet insoluble material and protein concentration was determined using Bradford protein assay. Equal amounts of protein (5 μg for cells and 20 μg for tissues) were prepared with loading buffer (1x Laemmli (Bio-Rad) with reducing agent (5% β-mercaptoethanol for cells and 100 mM DTT for tissues). Samples were heated at 37°C for 30 minutes and separated on a 20% SDS-PAGE. The gels were subsequently transferred to a PVDF membrane (Millipore), blocked in 5% milk in Tris-buffered saline/0.1% Tween-20 (TBS-T) and incubated with anti-sheep ESGP antibody (1 mg/μl;R&D Systems) overnight at 4°C. Membranes were then washed with TBS-T and incubated with Alexa-Fluor 647 donkey anti-sheep secondary antibody (1:5,000; Invitrogen). Bands were visualized using the Odyssey^®^ infrared detection system (LI-COR Biosciences). GAPDH (1:5,000; Millipore) was used as a loading control.

### Subcellular fractionation

C2C12 cells were harvested on day 2 of differentiation in ice cold hypotonic buffer (10 mM Tris-HCl pH 8, 2 mM EDTA) and lysed using a dounce homogenizer. Lysates were then centrifuged at 800 × g for 5 mintues at 4°C to separate nuclei and cell debris. That supernatant was then centrifuged at 5000 × g for 10 minutes to pellet mitochondria and ER. ER and heavy vesicles were further pelleted through centrifugation at 17,000 × g for 10 minutes. Finally, plasma membrane, light vesicles, and organelles were pelleted at 100,000 × g for 20 minutes and the supernatant from this spin was collected as the cytosolic fraction. All pellets were resuspended in lysis buffer (50 mM Tris-HCl, pH 6.8, 1mM EDTA, 2% SDS) at volumes equal to the supernatant. 8 μl of each fraction was separated by SDS-PAGE and analyzed for presence of myomerger, caveolin-3, and tubulin. Caveolin-3 antibody (BD Transduction Laboratories #610421) was used at 1:6700 and tubulin (Santa Cruz #SC-8035) at 1:50.

### Immunocytochemistry

Cultured cells were rinsed with PBS and fixed in 4% paraformaldehyde (PFA)/PBS for 15 minutes at room temperature. Cells were subsequently permeabilized and blocked in 0.01% Triton X-100/5% donkey serum/PBS for one hour at room temperature. Primary antibody diluted in permeabilization/blocking buffer was incubated overnight. Cells were then washed with PBS and incubated with secondary Alexa-Fluor antibodies (1:250) for 1 hour. A myomaker antibody custom antibody was generated through YenZym Antibodies LLC. Rabbits were immunized with amino acids #137-152 of mouse myomaker (MKEKKGLYPDKSIYTQ) after conjugation to KLH.

We used antigen-specific affinity purified products at a concentration of 4.3 μg/mL for immunostaining. Esgp (myomerger) antibody was used at a concentration of 1 μg/mL. Anti-mouse myosin (my32, MA5-11748, ThermoFisher Scientific) antibody was used at 1:100. Hoechst 33342 solution (ThermoFisher Scientific) was used to stain nuclei. Cells were imaged using Nikon A1R+ confocal on a FN1 microscope (35 mm dishes) or Nikon A1R confocal on Eclipse T1 inverted microscope (Ibidi slides).

### Histology and immunohistochemistry

For cryosections, embryos were dissected, fixed in 4% PFA/PBS overnight at 4°C, washed in PBS, incubated in 30% sucrose/PBS overnight and then in 1:1 mix of optimal cutting temperature (O.C.T.) formulation and 30% sucrose prior to embedding in O.C.T˙. Sections were cut at 10 μm and then permeabilized with 0.2% Triton X-100/PBS, blocked with 1% BSA/1% heat inactivated goat serum/0.025% Tween20/PBS and incubated with primary antibody overnight. Anti-mouse myosin (my32, MA5-11748, ThermoFisher Scientific) antibody was used at 1:100, whereas myogenin (F5D, Developmental Hybridomas) was used at a concentration of 2.56 μg/mL. Secondary goat anti-mouse IgG1-488 Alexa-Fluor antibody (Invitrogen) was incubated at a dilution of 1:250 for 1 hour. Slides were mounted with VectaShield containing DAPI (Vector Laboratories) and visualized using Nikon A1R confocal on Eclipse T1 inverted microscope. Images were analyzed with Fiji.

### Statistical analysis

For quantitation of cell fusion in Fig. 1b, cells with 3 or more nuclei were considered syncytial cells. The number of nuclei in syncytial cells and total number of nuclei were manually counted. In Fig. 1c, the number of myosin^+^ myotubes (myosin structures with 3 or more nuclei) and GFP^+^ myosin^+^ myotubes were manually counted. To quantify fusion between myomaker^+^ myomerger^+^ GFP^+^ fibroblasts with either myomaker^+^ NLS-Tom^+^ or myomerger^+^ NLS-Tom^+^ fibroblasts (Extended Data Fig. 3), we calculated the percentage of GFP+ NLS-Tom^+^ syncytial cells. The differentiation index (Fig. 3c) was calculated as the percentage of nuclei in myosin^+^ cells, and the fusion index (Fig. 3d) as the percentage of myosin^+^ cells with the indicated number of nuclei. Quantitative data sets are presented as means ± SEM. For each quantitiation, at least 3 independent experiments were performed in duplicate and 4-6 fields were randomly chosen for imaging. Histological analysis of embryos was performed on 3-4 embryos per genotype per time point. Multiple histological levels within each muscle were examined. The data were analyzed using an unpaired Student’s t-test with GraphPad Prism 6 software. A value of *P* < 0.05 was considered statistically significant.

## Acknowledgements

We thank the Transgenic Animal and Genome Editing Core and the Confocal Imaging Core at the Cincinnati Children’s Hospital Medical Center. This work was supported by grants to D.P.M. from the Cincinnati Children’s Hospital Research Foundation, National Institutes of Health (R01AR068286), and Pew Charitable Trusts.

## Author Contributions

D.P.M and M.E.Q. designed and performed experiments, and analyzed the data. Q.G., M.K., D.G.G., M.J.P., and V.K. performed experiments. D.P.M. wrote the manuscript with assistance from all authors.

## Author Information

The authors declare no competing financial interests. Correspondence and requests for materials should be addressed to D.P.M.

---

**Extended Data Figure 1.**
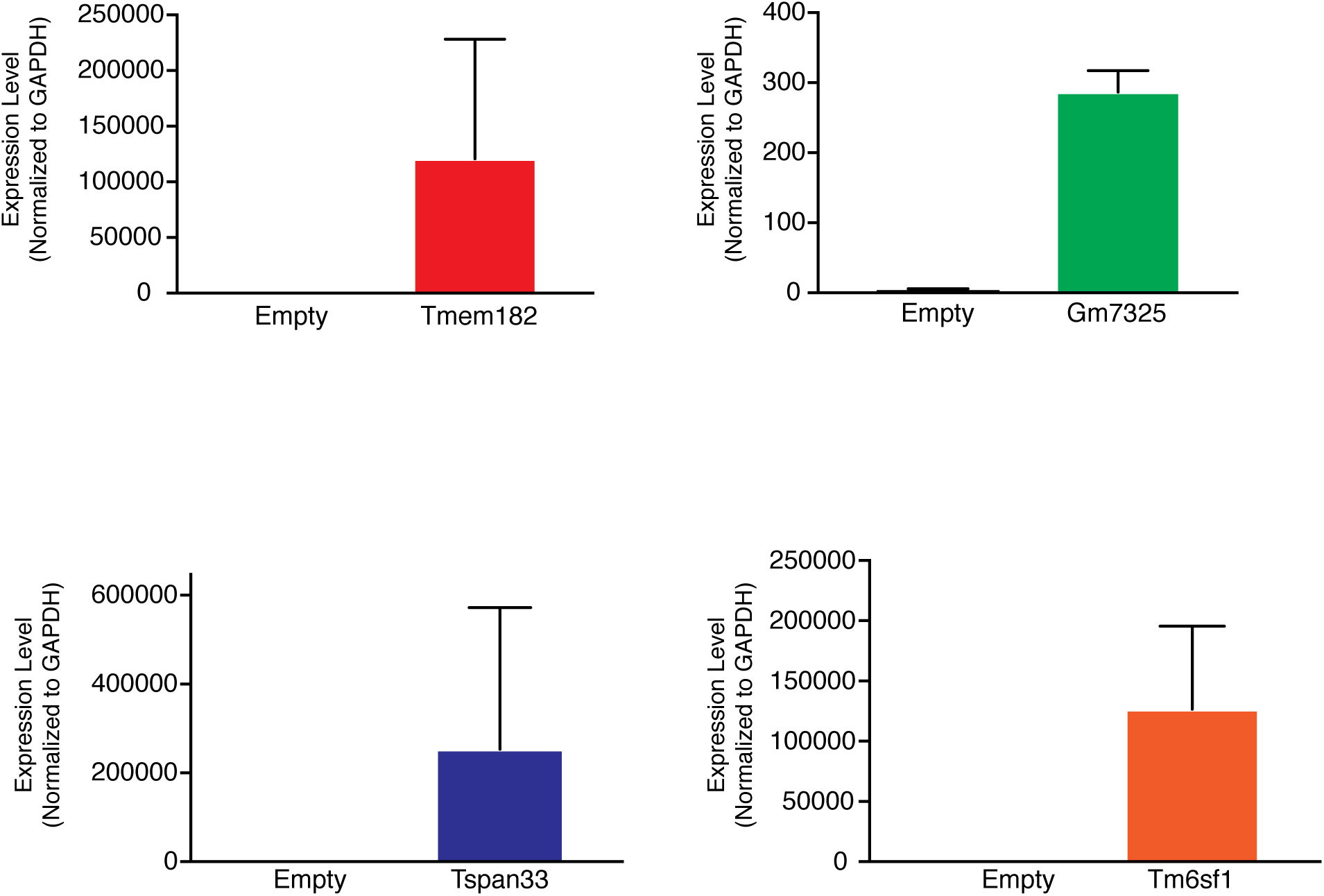
Expression of MyoD-regulated genes in myomaker^+^ fibroblasts. qRT-PCR analysis for the indicated genes 72 hours after expression in fibroblasts. For *Gm7325*, we used primers specific for the long transcript.

**Extended Data Figure 2.**
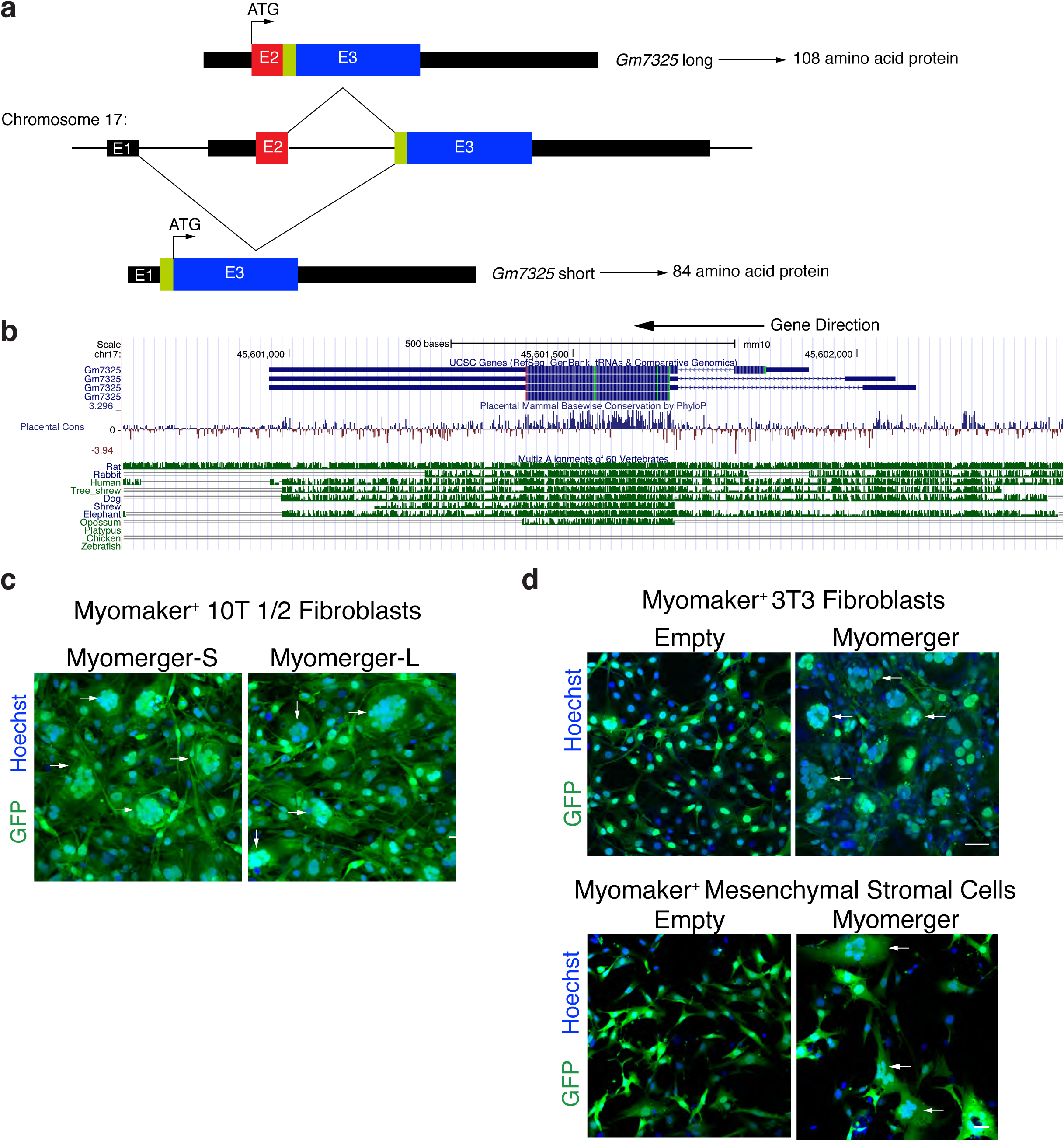
Genomic organization of murine *Gm7325* and the ability of both variants to induce fusion. **a,** Diagram showing the Gm7325 locus on chromosome 17. The short transcript is generated by splicing of exon 1 (non-coding) with exon 3, leading to an 84 amino acid protein. The long transcript is produced by splicing of exon 2 with exon 3 and results in a 108 amino acid protein. **b**, UCSC genome browser track showing multiple transcripts and conservation across vertebrate species. The short transcript is highly conserved in multiple species, including human, but not present in zebrafish. The upstream exon that produces the longer transcript is not highly conserved. Note that this annotation displays the gene on the reverse strand. **c**, The short (S) or long (L) myomerger transcripts were expressed in myomaker^+^ 10T ½ fibroblasts and both induced fusion (*n*=3). **d**, Myomerger also induced fusion of myomaker^+^ NIH/3T3 fibroblasts (*n*=3) and myomaker^+^ mesenchymal stromal cells (*n*=3). Arrows indicate fusion. Scale bars, 50 μm.

**Extended Data Figure 3.**
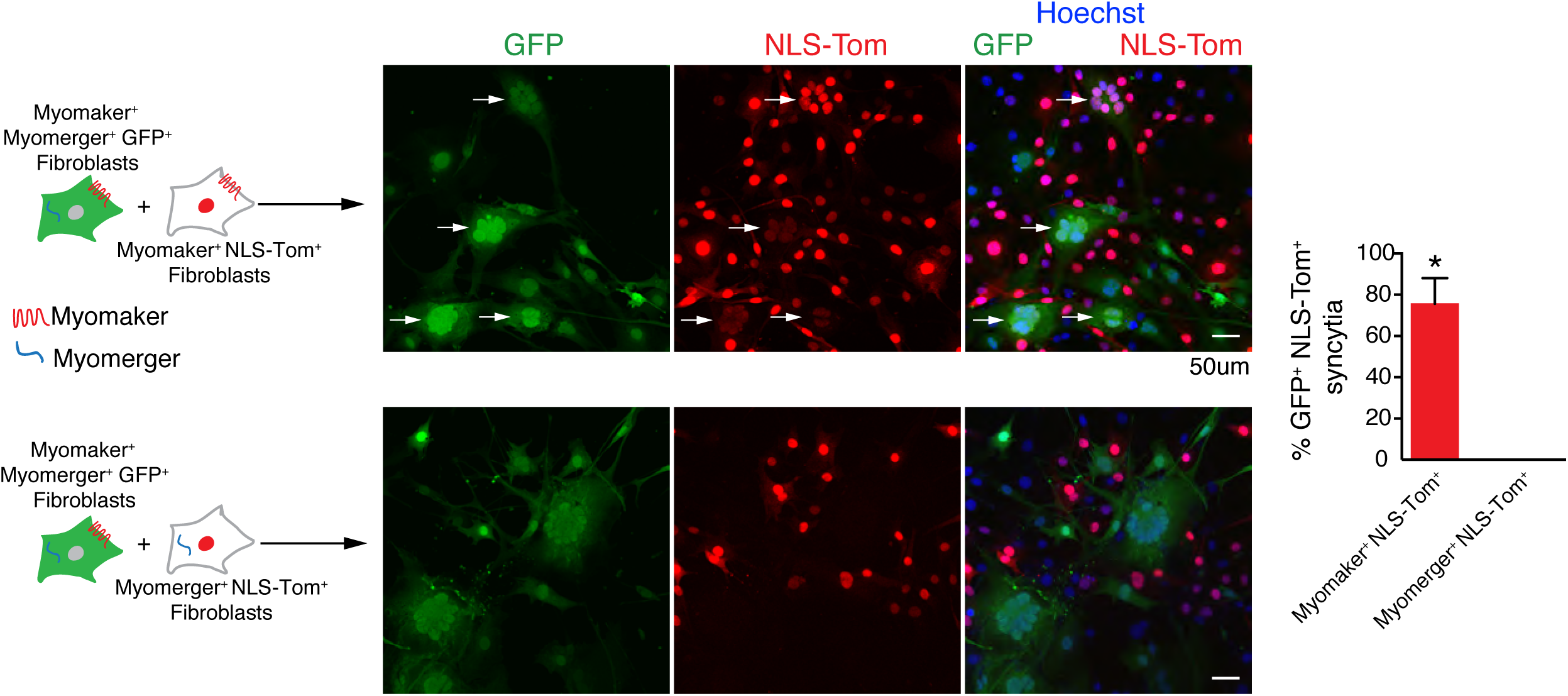
Efficient fusion requires myomaker expression in both fusing cells but myomerger in one fusing cell. Diagram showing the cell mixing approach to assess fusion between the populations of fibroblasts. Co-localization of GFP and NLS-TdTomato (NLS-Tom) in the nucleus represents fusion (arrows). Representative images demonstrate fusion of myomaker^+^ myomerger^+^ GFP^+^ fibroblasts with myomaker^+^ NLS-Tom^+^ fibroblasts but not myomerger^+^ NLS-Tom^+^ fibroblasts. The percent of GFP^+^ NLS-Tom^+^ syncytia (*n*=3). Data are presented as mean ± SEM. \**P*<0.05 compared to myomerger^+^ NLS-Tom^+^ fibroblasts. Scale bars, 50 μm.

**Extended Data Figure 4.**
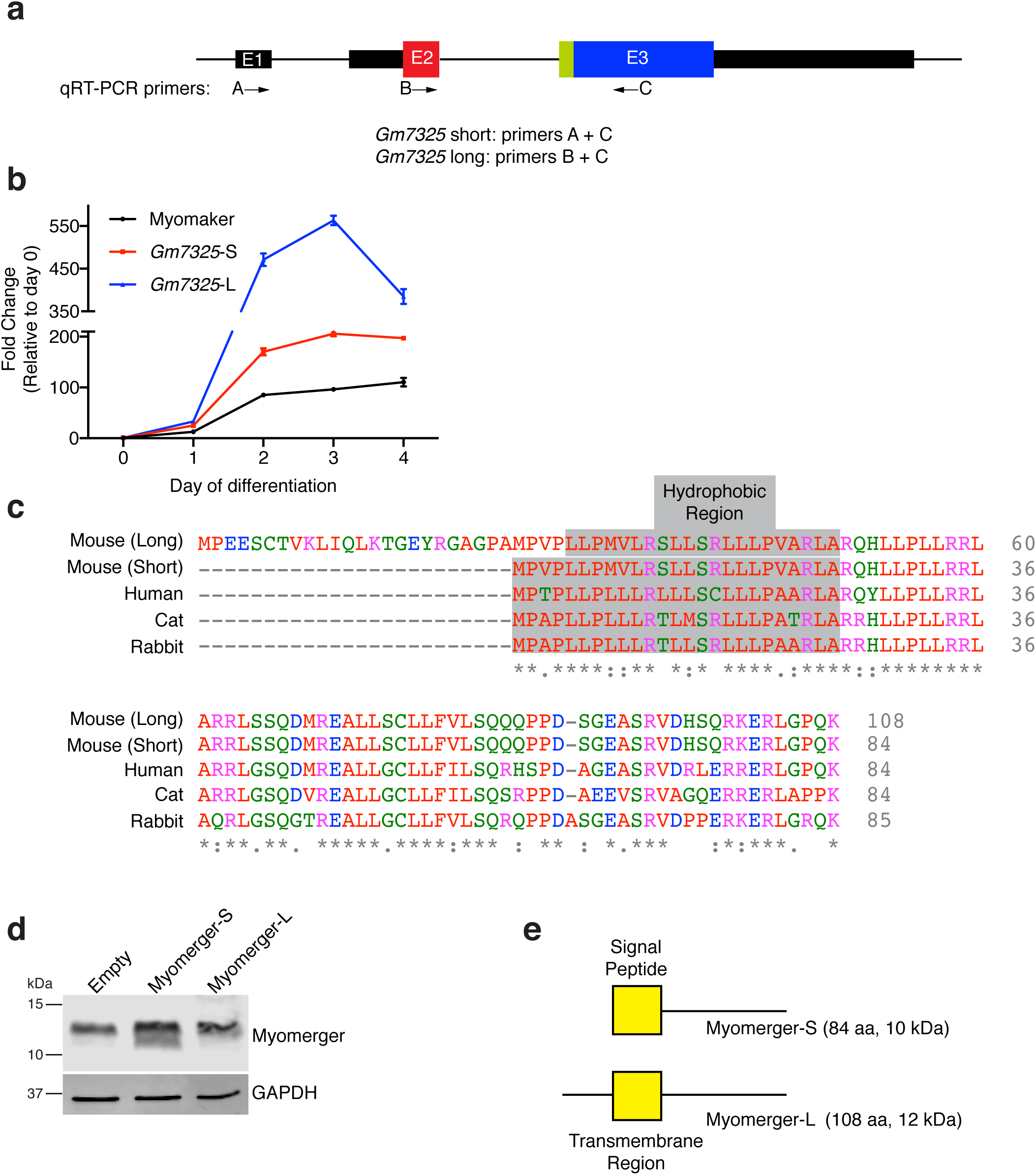
Design of qRT-PCR primers and comparison of myomerger protein variants. **a**, Schematic showing the location of primers to distinguish short and long transcripts. **b**, qRT-PCR for *Gm7325* transcript variants and myomaker in C2C12 cells on the indicated days of differentiation (*n*=*3* for each time point). **c**, Sequence alignment of both mouse myomerger protein products with multiple mammalian orthologs using Clustal Omega. A potential hydrophobic region is highlighted in gray. **d**, Immunoblotting from C2C12 cells infected with either empty, myomerger-short (S), myomerger-long (L) on day 2 of differentiation. Myomerger migrates as a single band around 12 kDa when endogenously produced (empty). Over-expression of myomerger-S leads to an increase in the endogenous band and a lower band is also detected suggesting that myomerger transcripts may be subjected to intricate mRNA processing or post-translational modifications. **e**, Graphic showing the regions of myomerger-S and myomerger-L as predicted by SignalP and Phobius.

**Extended Data Figure 5.**
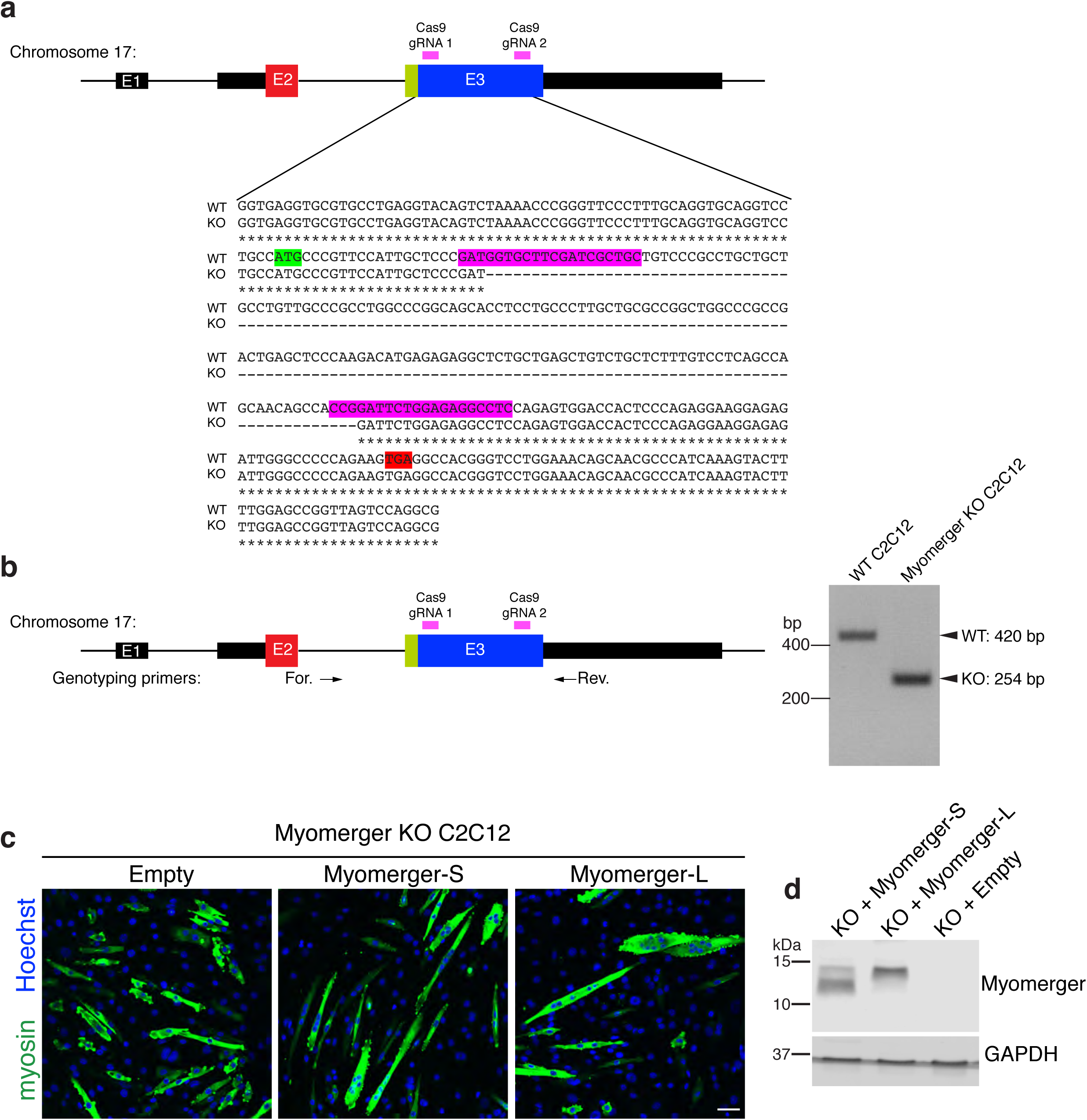
CRISPR/Cas9 disruption of the *Gm7325* locus. **a**, Schematic showing the Gm7325 locus and targeting of sgRNAs. **b**, Genotyping strategy for myomerger KO C2C12 cells. WT and KO PCR products were sequenced and the result is shown below the diagram in (**a**). The use of two sgRNAs results in reproducible cut sites leading to a 166 base pair deletion in both C2C12 cells and mice. The translational start site (ATG, green) for myomerger-S and stop site (TGA, red) for both myomerger-S and myomerger-L are noted. **c**, Myomerger KO C2C12 cells were infected with either empty, myomerger-S, or myomerger-L and induced to differentiate. Both myomerger-S and myomerger-L rescued the lack of fusion in myomerger KO cells. **d**, Immunoblotting for myomerger shows appropriate expression after transduction of myomerger KO cells. Scale bars, 50 μm.

**Extended Data Figure 6.**
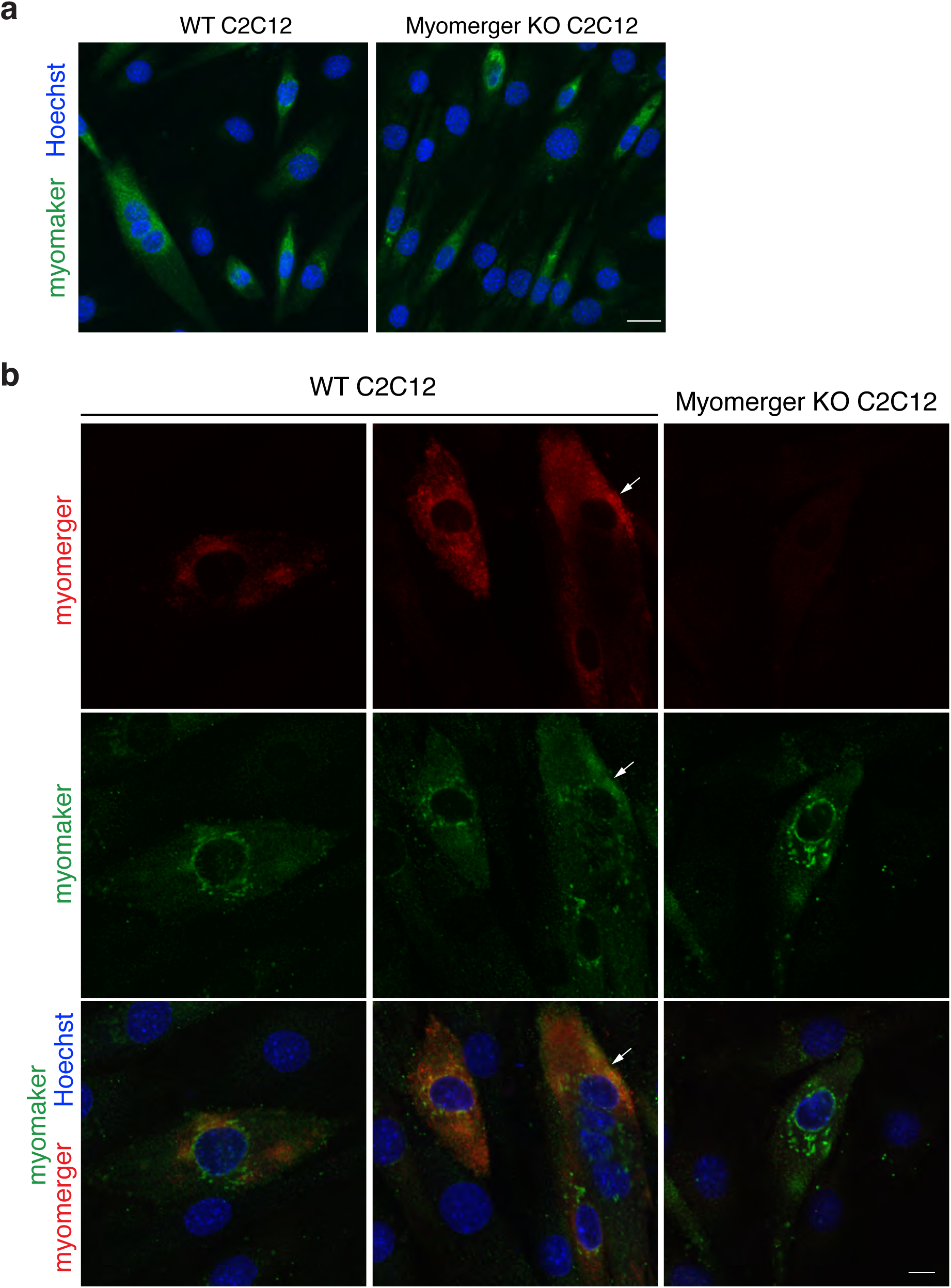
Analysis of myomaker and myomerger co-localization. **a**, Representative immunofluorescence images from WT and myomerger KO C2C12 cells on day 2 of differentiation indicating that loss of myomerger does not alter myomaker expression or localization. **b**, Immunofluorescence for myomerger and myomaker on the indicated cells on day 2 of differentiation. These two fusion proteins exhibit different patterns, although modest co-localization (arrows) was observed. The functional significance of this potential co-localization is not currently known. Scale bars, 10 μm (**a**), 5 μm (**b**).

**Extended Data Figure 7.**
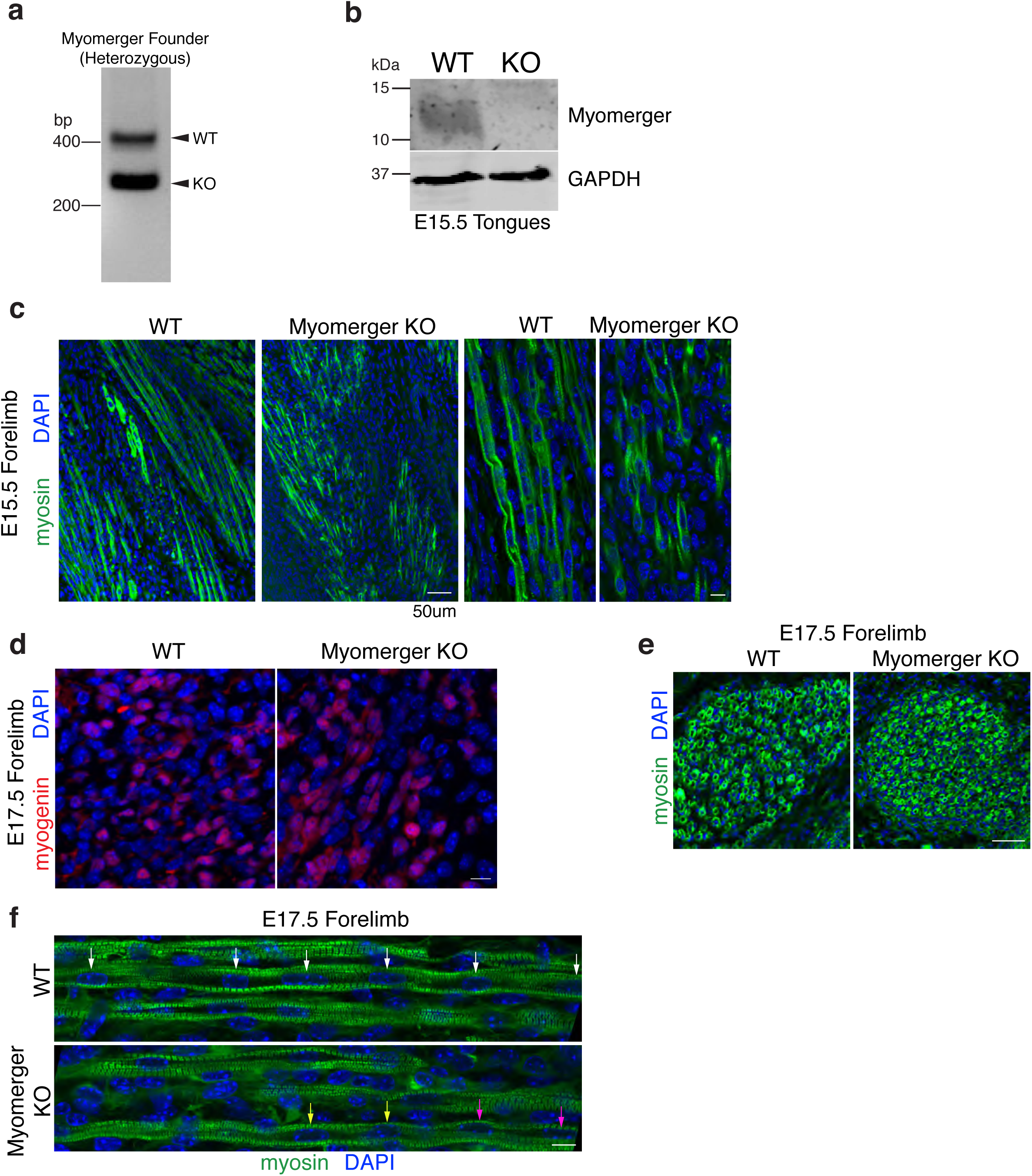
Examination of myomerger KO muscle. **a**, Genotyping of the one founder harboring the *Gm7325* mutation generated through Cas9-mutagensis. **b**, Immunoblotting on tongue lysates from WT and myomerger KO mice showing lack of myomerger in KO samples. GAPDH was used as a loading control. **c**, E15.5 forelimbs (*n*=3) immunostained with a myosin antibody demonstrates that myomerger KO myoblasts differentiate but are unable to fuse. **d-e**, E17.5 forelimbs from WT and myomerger KO mice were evaluated for myogenin and myosin expression, and multi-nucleation. Arrows of same color in (**f**) show nuclei within one myofiber. We observed myocytes in myomerger KO samples that contained two nuclei (arrows). Analysis of z-stack images (not shown) indicated that the nuclei labeled by the yellow and pink arrows are not within the same myofiber. Scale bars, 50 μm (**c**, left panels, **e**), 10 μm (**c**, right panels, **d, f**).

